# The *Chlamydia trachomatis* type III secreted effector protein CteG induces centrosome amplification through interactions with centrin-2

**DOI:** 10.1101/2022.06.23.496711

**Authors:** Brianna Steiert, Carolina M. Icardi, Robert Faris, Aloysius J. Klingelhutz, Peter M. Yau, Mary M. Weber

## Abstract

The centrosome is the main microtubule organizing center of the cell and is crucial for mitotic spindle assembly, chromosome segregation, and cell division. Centrosome duplication is tightly controlled, yet several pathogens, most notably oncogenic viruses, perturb this process leading to increased centrosome numbers. Infection by the obligate intracellular pathogen *Chlamydia trachomatis* (*C.t.*) correlates with blocked cytokinesis, supernumerary centrosomes, and multipolar spindles; however, the mechanisms behind how *C.t.* induces these cellular abnormalities from the confines of its inclusion, remain largely unknown. Here we show that the type III secreted effector protein, CteG, binds to centrin-2 (CETN2), a key structural component of centrosomes and regulator of centriole duplication. This interaction requires a functional calcium binding EF hand 4 of CETN2, which is recognized via the C-terminus of CteG. Significantly, we show that deletion of CteG, or knockdown of CETN2, significantly impairs chlamydia’s ability to induce centrosome amplification. Uniquely, we have identified the first bacterial effector to target centrins, crucial regulators of the eukaryotic cell cycle. These findings have not only allowed us to begin addressing how *C.t.* induces gross cellular abnormalities during infection, but also indicate that obligate intracellular bacteria may contribute to cellular transformation events that negatively impact host physiology even when the pathogen is long removed. Understanding the consequences of CteG-CETN2 interactions, its impact on centrosome amplification, and the long-term effect this has on host cells could explain why chlamydial infection leads to an increased risk of cervical or ovarian cancer.

**Significance Statement:** The presence of more than two centrosomes is a hallmark of many types of cancer, including cervical and ovarian cancers of which *Chlamydia trachomatis* (*C.t.*) infection is a significant risk factor. Despite the importance of this problem, how *C.t.* orchestrates these drastic changes in the host cell remains poorly understood. Here, we describe how *C.t.* uses a single effector protein, CteG, to drive centrosome amplification via manipulation of a key regulator of centriole duplication, centrin-2. This work begins to define how *C.t.* induces centrosome amplification to promote its replication while potentially contributing to devastating long-term negative consequences for normal host physiology. Further it may help elucidate why chlamydial infection leads to an increased cancer risk.

## Introduction

The centrosome is the main microtubule organizing center (MTOC) of the cell and is involved in mitotic spindle assembly, chromosome segregation, cell division, microtubule structure, and cell shape (1). The centrosome is comprised of two barrel-shaped centrioles that are embedded in a matrix of proteins known as the pericentriolar material. Centrosomes duplicate only once per cycle, initiating the process at the G1/S phase transition and completing this process prior to entry into mitosis (2). Given the intimate link between cell cycle progression and centrosome duplication, there is increasing support for the notion that the centrosome itself is a key regulator of the cell cycle (3).

Centrosome abnormalities are hallmarks of numerous types of human cancers and correlate with tumorigenesis and poor patient outcomes (4). Centrosome amplification can be caused by cell-cell fusion, dysregulation of centrosome duplication, or cytokinesis defects (5). Centrosome amplification leads to increased genomic instability, which can increase merotelic attachment of kinetochores, resulting in aneuploidy and chromosomal instability (6), both of which are additional hallmarks of cancer. While centrosome amplification has been associated with these other hallmarks of cancer, centrosome amplification alone has also been shown to be sufficient to cause tumorigenesis in flies and mammals (7, 8). Typically, increased centrosome number alters mitotic spindle formation, leading to multipolar spindles, which can support tumorigenesis by promoting merotelic attachments and chromosome mis-segregation (9, 10). Division in cells with multipolar spindles can be deleterious, but cancer cells overcome this by clustering extra centrosomes to achieve bipolar mitosis (5, 11). Viruses linked to increased cancer risk, such as human papillomavirus (HPV) and Epstein-Barr virus (EBV), have been shown to similarly induce centrosome abnormalities. Cervical cancers associated with high-risk HPV infection are characterized by multipolar spindles, which is linked to abnormal centrosome number (12). The HPV oncoprotein E7 induces centrosome amplification by targeting centriole duplication, which can lead to centrosome accumulation and ultimately causes genomic instability. Similarly, EBV infection leads to overproduction of centrosomes through its BNRF1 protein (13).

*Chlamydia trachomatis* (*C.t.*) is an obligate intracellular bacteria that is the etiological agent of multiple human diseases (14). Importantly, current or prior chlamydia infection has been associated with an increased risk for development of ovarian and cervical cancer (15, 16). Chlamydia is known to cause host cell transformation and it has been speculated that *C.t.*-induced changes to the host cell linger after clearance of infection (17), potentially explaining why chlamydial infection increases the risk of developing certain types of cancers. Early during infection, *C.t.* traffics along microtubules to the MTOC of the cell to establish its intracellular niche, termed the inclusion. Here it maintains a close association with the MTOC/centrosomes (17). Studies have shown that chlamydia infection leads to supernumerary centrosomes, mitotic spindle defects, multinucleation, aneuploidy, and blocked cytokinesis (17–21). In *C.t.* infection models, centrosome amplification has been attributed to both cytokinesis defects and dysregulation of the centrosome duplication machinery (18, 19, 22). While the initial observations that *C.t.* infection induces gross host cellular abnormalities were made over 15 years ago, how *C.t.* orchestrates these cellular changes from the confines of its inclusion remains largely unknown.

As an obligate intracellular pathogen, *C.t.* must establish a niche within a host to proliferate and cause disease. Essential to this intracellular lifestyle is the secretion of over 100 effector proteins, which are delivered through its type III secretion system (T3SS) (23, 24). These effector proteins have been shown to play roles in invasion, nutrient acquisition, and immune evasion, but the function of most remains unknown (23, 24). The *<u>C</u>hlamydia trachomatis* <u>e</u>ffector associated with the <u>G</u>olgi (CteG) is a T3SS effector that was previously shown to localize to the Golgi or plasma membrane depending on the stage of the infection cycle (25). When expressed in yeast, CteG causes a vacuolar protein sorting defect (25), however the molecular function of CteG remains unknown.

In this study, we investigated the role of CteG during *C.t.* infection. We detected a novel interaction between CteG and the host protein, centrin-2 (CETN2), and further demonstrate that binding requires the C-termini of both proteins. Significantly, our results indicate that CteG is necessary for centrosome amplification but is dispensable for multinucleation and centrosome positioning at the inclusion. Intriguingly, while CteG is dispensable for growth in immortalized cell lines that possess supernumerary centrosomes, the absence of CteG impairs chlamydia’s ability to replicate efficiently in primary cervical cells. Thus, our new data indicates that CteG is a primary contributor to *C.t.* induced centrosome amplification via manipulation of centrin-2 (CETN2) and this interaction is important for chlamydial growth.

## Results

### CteG toxicity in yeast is suppressed by overexpression of the Anaphase Promoting Complex Subunit 2

Previous studies demonstrated that expression of CteG in yeast resulted in a vacuolar protein sorting defect (25); however, the precise mechanism and function remains unknown. To further dissect the function of CteG, we exploited yeast genetics to identify the host pathway(s) targeted by this effector protein. In line with previous observations (26), we demonstrate that when CteG is overexpressed in yeast, a clear toxic phenotype is observed (Fig. 1A), suggesting that CteG perturbs an essential host pathway. To identify the target pathway(s), we employed a yeast suppressor screen (27–29). Introduction of a yeast genomic library into the CteG-expressing yeast strain yielded 69 putative suppressor colonies, of which 25 markedly reduced CteG-induced toxicity. Sequencing revealed that 18 of these clones harbored the ORF APC2. When expressed independently, the anaphase promoting complex subunit 2 (APC2) was sufficient to suppress CteG toxicity in yeast, but did not suppress the toxicity of TmeA (Fig. 1A), a cT3SS effector previously shown to target N-WASP (30, 31). APC2 is a subunit of the anaphase promoting complex (APC), a ubiquitin ligase involved in the degradation of cyclins to promote the progression of the cell cycle (11). The yeast suppressor screen provides putative targeted pathway(s), not direct interacting partners, so these data suggest that CteG targets pathway(s) involved in the host cell cycle.

**Fig. 1.**
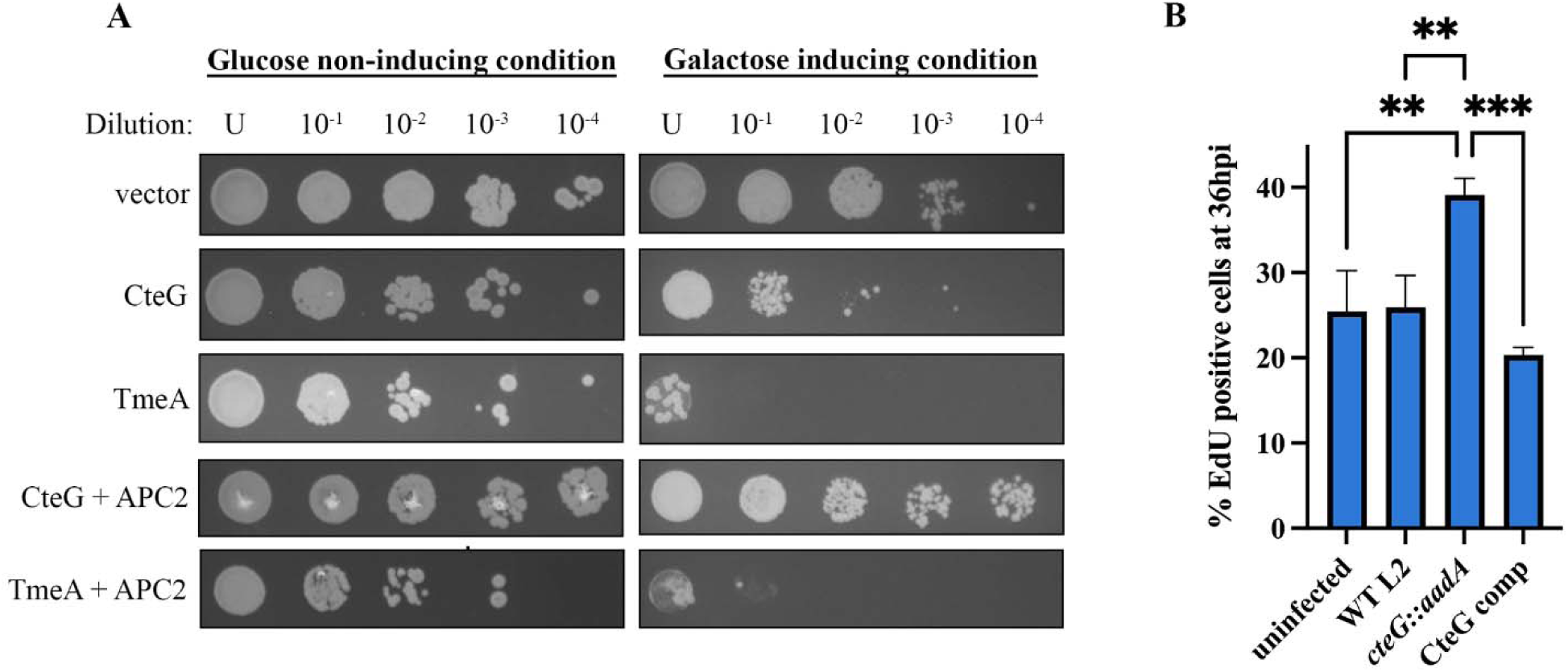
CteG toxicity in yeast is suppressed by APC2 and contributes to altered cell cycle progression. (*A*) *C.t.* effectors and APC2 were placed under the control of galactose inducible promoters and were serially diluted and spotted onto glucose- or galactose-containing media. (*B*) Quantification of EdU positive HeLa cells 36 hours post-infection. Significance was determined using one-way ANOVA followed by Tukey’s multiple comparisons test. Error bars are SD, ** P< 0.01, *** P<0.001. Data are representative of 2 replicates.

### Cells infected with a CteG null mutant show altered cell cycle progression

To determine if CteG is playing a role in perturbing the host cell cycle, we labeled cells with EdU at 36 hours post-infection. Cells were either left uninfected or infected with WT L2, CteG null mutant (*cteG::aadA*) (25), or *cteG::aadA* complemented with pBomb4-tet-CteG-FLAG (CteG comp.). At 36 hours post-infection, we found a higher percentage of *cteG::aadA* infected cells were in S-phase, indicating that these cells are progressing faster through the host cell cycle compared to WT or CteG comp. infected cells (Fig. 1B). This suggests that CteG’s role within the host may influence progression of the host cell cycle, either directly, or downstream of the effects of its direct target.

### CteG binds to the host protein centrin-2

To identify the physiologically relevant target of CteG, we employed affinity purification-mass spectrometry (AP-MS). Expression of FLAG-tagged CteG was confirmed prior to AP-MS analysis by western blotting (Fig. S1). For analysis of MS data, we compared FLAG tagged CteG results to those of empty vector and only peptides unique to CteG were considered for further analysis. To further narrow our list of peptides, those with an average of 1 peptide count across all three replicates were removed, leaving 76 putative targets (Table S2). With an average peptide count of 9 and an average of 21 matches across three biological replicates, a fragment of centrin-2 (CETN2) was the second most abundant unique peptide (Table 1, Table S2). CETN2 is a key structural component of centrosomes and regulator of centriole duplication (32). Our yeast suppressor screen suggests a role for CteG in perturbing the host cell cycle, which may have centrosome-based checkpoints as centrosome defects lead to perturbations of the cell cycle (3). As CETN2 is a component of centrosomes, we sought to validate that CteG interacts with CETN2. We immunoprecipitated FLAG-tagged CteG from infected cells transfected with HA-tagged CETN2. Only FLAG tagged CteG pulled down HA-tagged CETN2. No interaction with vector or TmeA was noted (Fig. 2A). We further confirmed these findings using an anti-CETN2 antibody to probe for an interaction with endogenous CETN2 (Fig. S2). To determine whether this interaction is independent of other bacterial factors, we co-transfected cells with HA-tagged CETN2 and GFP-tagged CteG, TmeA, or empty vector control. Again, CteG uniquely co-immunoprecipitated with CETN2 (Fig. 2B). In addition, cells co-transfected with CETN2-dsRed and GFP-tagged CteG were observed to co-localize, as evident by a significant Pearson’s R value (Fig. 2C). No co-localization with the negative controls GFP or TmeA-GFP was noted. Collectively, our results indicate that CteG specifically binds to CETN2, and that this interaction does not require any additional bacterial factors. Significantly, CteG represents the first bacterial effector protein identified that targets centrin proteins.

**Fig. 2.**
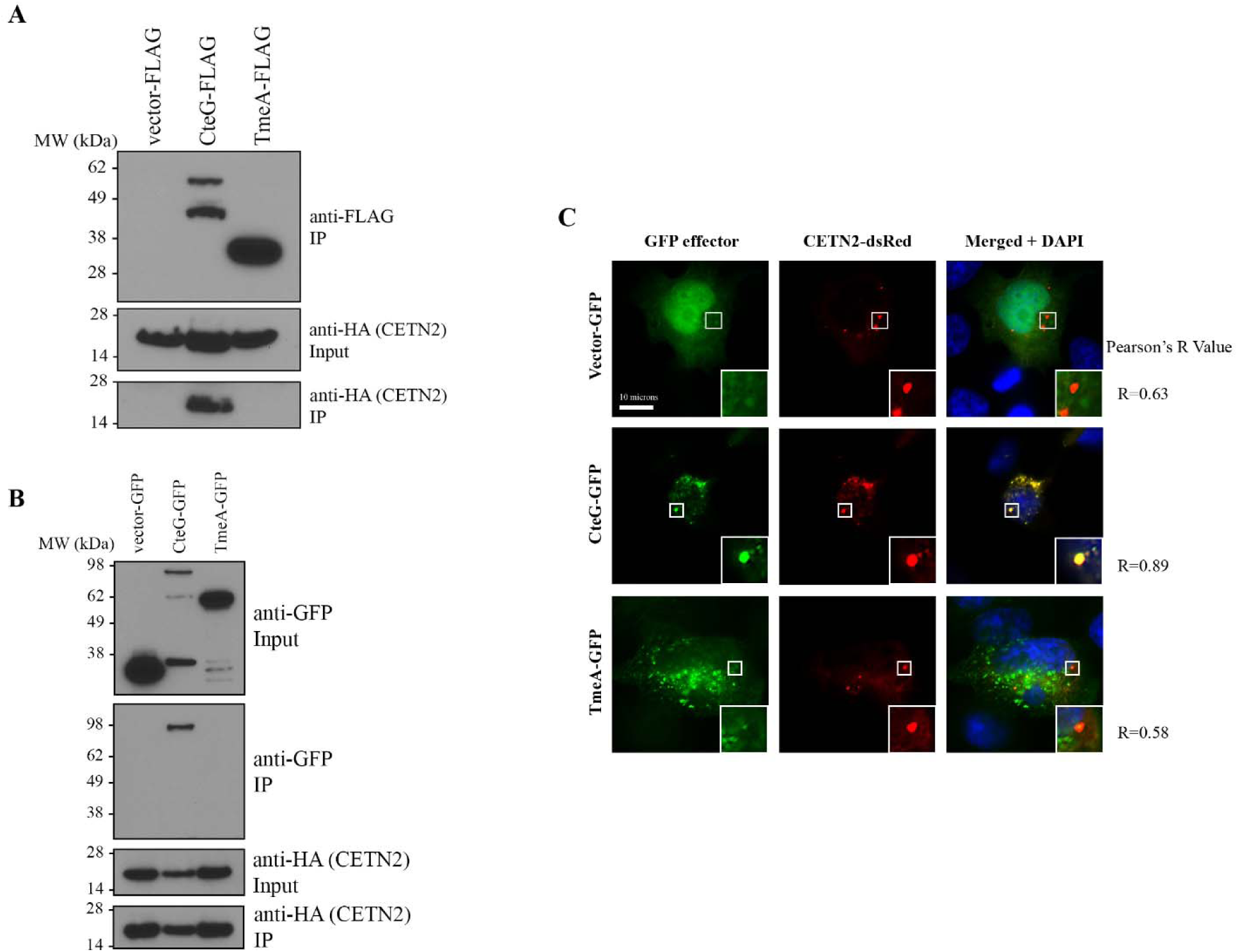
CteG interacts with CETN2. (*A*) Co-IP of *C.t.* expressing FLAG-tagged effectors from CETN2-HA transfected HeLa cells. (*B*) Co-IP of HA-tagged CETN2 from HeLa cells co-transfected with GFP-tagged effectors. (*C*) Immunofluorescence images of HeLa cells co-transfected with CETN2-dsRed (red) and empty vector-, CteG-, or TmeA-GFP (green). Nucleus is stained with DAPI (blue) (Scale bar, 10 microns). Data are representative of 2-3 replicates.

**Table 1.**
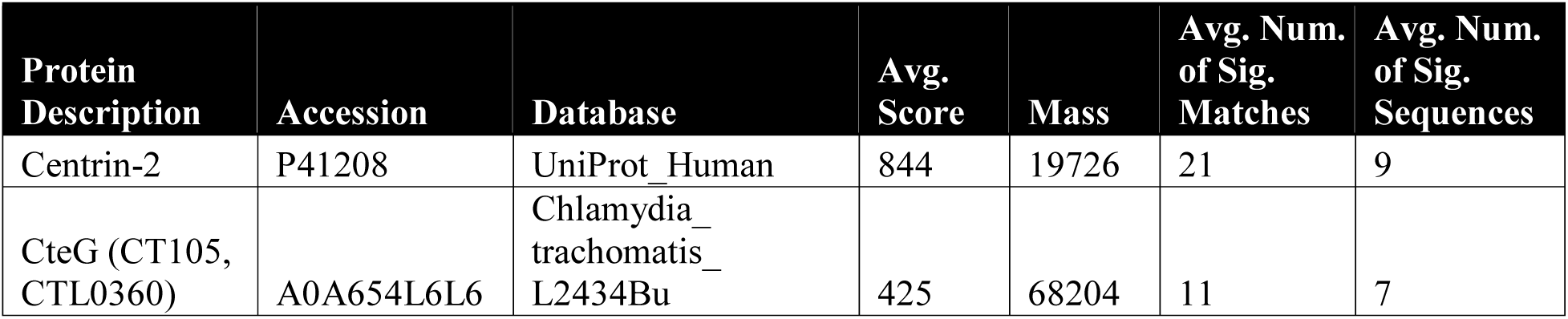
AP-MS peptides of interest for CteG.

### The C-terminal 23 amino acids of CETN2 are necessary for CteG binding

CETN2 has multiple phosphorylation sites and four EF-hand domains capable of calcium binding (33). These phosphorylation sites and EF hands are important for centrin localization and formation of centrin-containing structures at the MTOC (33, 34). To determine where CteG is binding CETN2, we made sequential (∼100 nucleotide/33 amino acid) truncations from the C- and N-termini of CETN2 (Fig. 3A). Truncations were cloned into pcDNA3.1+N-eGFP and transfected into HeLa cells, followed by infection with *C.t.* expressing FLAG-tagged CteG. Of these truncations, only the full length CETN2 bound to CteG, indicating the last 39 amino acids of CETN2 are important for binding (Fig. 3B, C). This C-terminal region of the protein includes EF hand 4, so further truncations were made before and after the calcium binding region of EF hand 4 (Fig. 3A). CteG co-immunoprecipitated with full length CETN2 and CETN2 truncated after the calcium binding domain of EF hand 4 (denoted CETN2 1-162) (Fig. 3C). This indicates that EF hand 4 is important for binding CteG. To determine the necessity of a viable EF hand, we made conserved mutations in EF hand 4 (**D**R**D**G**DG**-->**S**R**S**G**SA**) (Fig. 3D). This C-terminal region also contains a key phosphorylation site at serine 170, so we mutated this serine to alanine to prevent phosphorylation (denoted S170A CETN2) (Fig. 3D). Using a similar transfection/infection experiment, we show that CteG co-immunoprecipitated with full length CETN2, as well as S170A CETN2. No co-immunoprecipitation of the EF hand 4 domain mutant was noted (Fig. 3E). This indicates the importance of an intact EF hand 4 calcium binding domain for CteG-CETN2 interaction.

**Fig. 3.**
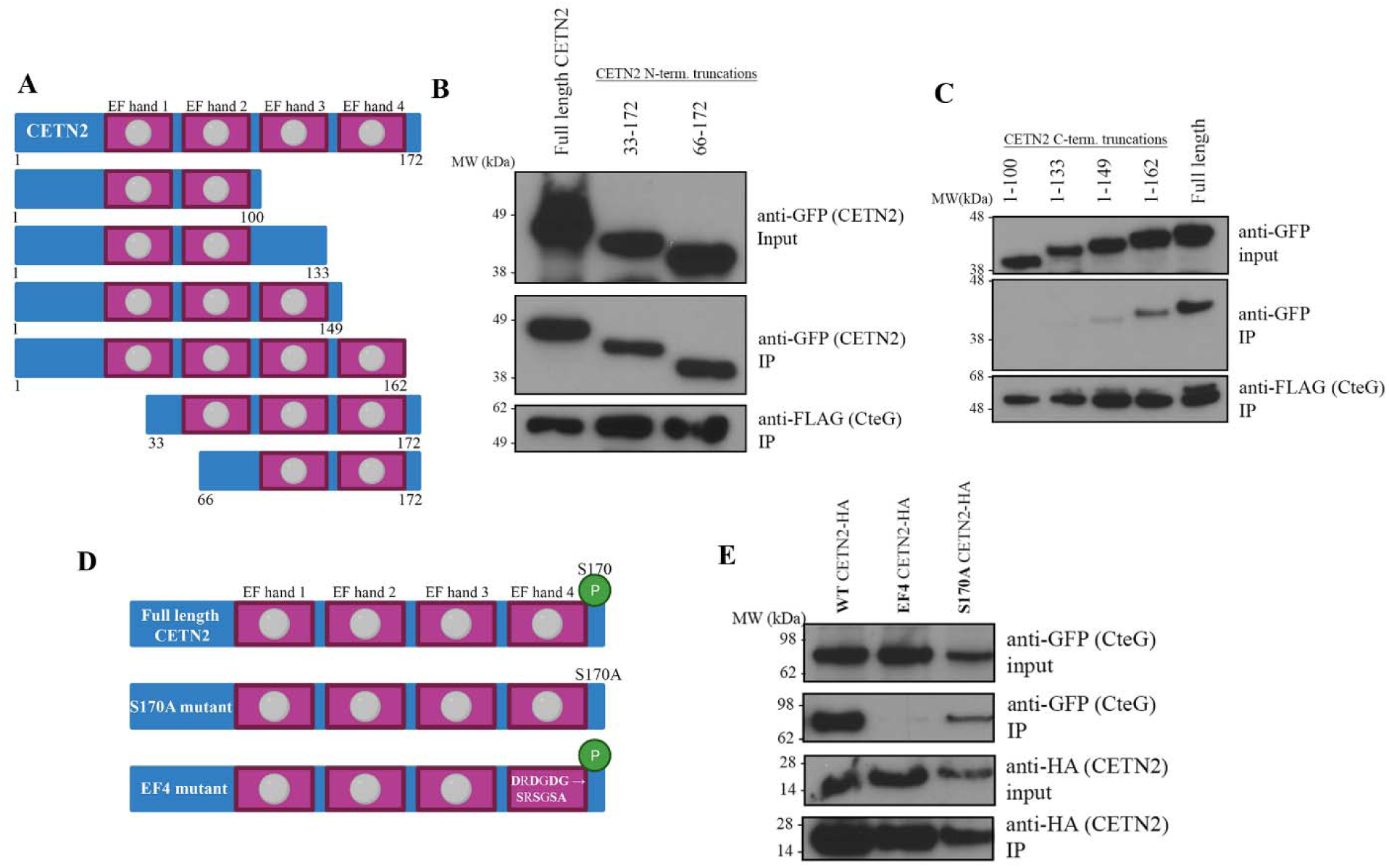
Intact C-terminus of CETN2 is needed for CteG-CETN2 interaction. (*A*) Schematic of CETN2 truncations made with full length CETN2 at the top. Pink boxes represent EF hand domains. Grey circles indicate calcium binding domains. (*B/C*) Co-IP of FLAG-tagged CteG expressing *C.t.* from HeLa cells transfected with N- or C-terminal truncations of CETN2. (*D*) Schematic of CETN2 mutations made to phosphorylation site S170 and the calcium binding domain of EF hand 4. Top figure shows intact calcium binding domains and S170 phosphorylation site (green circle). (*E*) Co-IP of HA-tagged CETN2 mutants from HeLa cells co-transfected with GFP-tagged CteG. Data are representative of 2-3 replicates.

### The C-terminus of CteG is necessary for CETN2 binding

To identify the regions of CteG that were necessary for this interaction, we made sequential C- and N-termini truncations and used co-transfection immunoprecipitations to determine regions necessary for binding CETN2. From these experiments, we found that the N-terminus is dispensable (Fig. S3). For C-termini truncations, we constructed FLAG-tagged C-terminal truncations of CteG and cloned them into the pB4-tet-mCherry plasmid (35) and *C.t.* was used to infect cells transfected with HA-tagged CETN2. Our data indicates the last 17 amino acids of CteG are important for CETN2 binding as only full-length CteG co-immunoprecipitated with CETN2 (Fig. 4A). To further confirm this, we transformed this truncated version of CteG into *S. cerevisiae* to determine if toxicity was lost without the C-terminus of CteG. Deletion of the last 17 amino acids of CteG resulted in loss of toxicity when overexpressed in yeast (Fig. 4B). Taken together, these experiments indicate the C-terminus of CteG is pertinent for binding to CETN2.

**Fig. 4.**
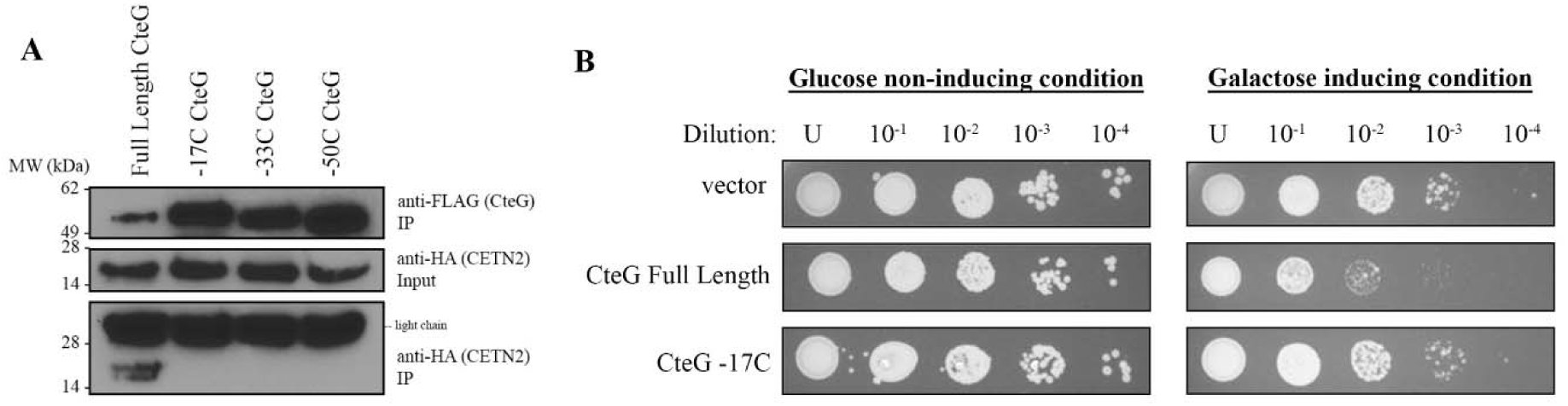
The C-terminus of CteG is necessary for CteG-CETN2 interaction. (*A*) Co-IP of FLAG-tagged *C.t.* expressing CteG truncations in HeLa cells transfected with CETN2-HA. (*B*) Yeast transformed with empty vector, full length CteG, and CteG -17C under galactose inducible promoters were diluted and spotted onto glucose- or galactose-containing media to assess toxicity. Data are representative of 2-3 replicates.

### Chlamydia amplifies centrosomes in a CteG-dependent manner

It is well established that chlamydia infection can cause gross host cellular abnormalities, including centrosome amplification (17–19), but the mechanisms behind this are unknown. Since CteG interacts with a key structural component of the centrosome important for centriole duplication, we sought to determine if CteG is essential for centrosome amplification during *C.t.* infection. A significant decrease in the percentage of cells with >2 centrosomes was noted in cells infected with the *cteG::aadA* relative to WT L2, CT144*::bla,* and CteG comp. (Fig. 5A, B) However, the presence of supernumerary centrosomes was still elevated compared to uninfected cells, indicating that CteG may not be the sole contributor to centrosome amplification during infection. Significantly, this same statistically significant decrease in centrosome amplification was observed between WT L2 and *cteG::aadA* infected primary cervical cells (Fig. 5C), further confirming the role of CteG in centrosome amplification. Host cellular abnormalities commonly associated with *C.t.* infection, such as multinucleation and altered centrosome positioning occurred independently of CteG expression, emphasizing the specific role of CteG in centrosome amplification (Fig. S4).

**Fig. 5.**
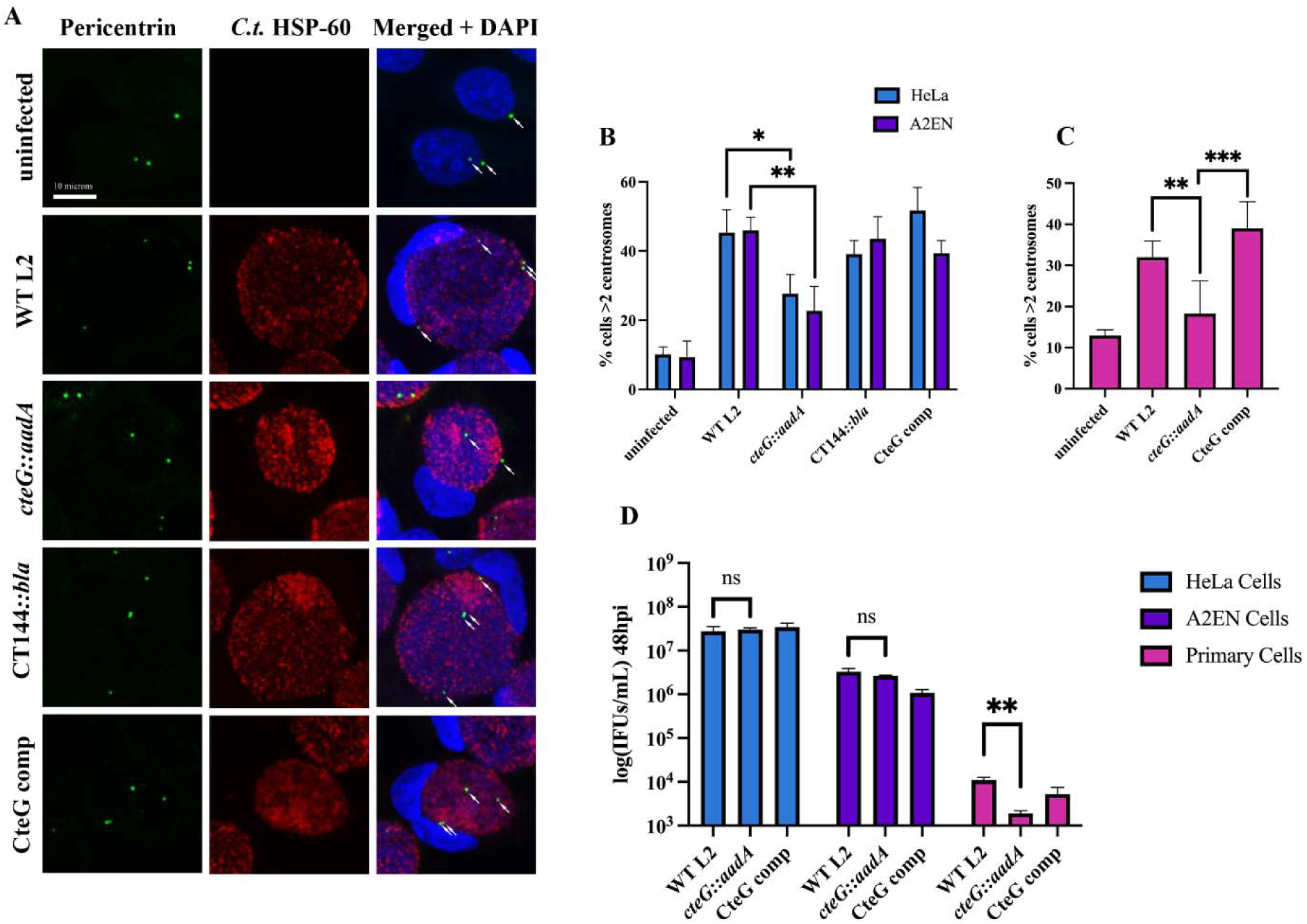
Centrosomes are amplified in a CteG dependent manner. (*A*) Representative images of A2EN cells infected with WT L2, *cteG::aadA*, CT144*::bla*, or CteG comp for 36h. Cells were stained with *C.t.* HSP-60 (red), pericentrin (green), and DAPI (blue). White arrows indicate centrosomes (Scale bar, 10 microns). (*B, C*) Quantification of cells with supernumerary centrosomes (>2) at 36 hours post infection in A2EN and HeLa cells (*B*) or primary cervical cells (*C)*. (*D*) Quantification of infectious progenies at 48 hours post-infection normalized to WT L2 IFUs at 0 hours. (*B-D*) Error bars are SD, * P< 0.05, ** P< 0.01, *** P<0.001. Significance was determined using one-way ANOVA followed by Tukey’s multiple comparisons test. Data are representative of 2-3 replicates.

As CteG is important for centrosome amplification, we next sought to determine whether CteG, and by extension supernumerary centrosomes, are important for chlamydial replication. While no growth defect was noted in HeLa and A2EN cells, a significant decrease in infectious progeny from *cteG::aadA* was noted compared to WT L2 and CteG comp infected cells (Fig. 5D). Collectively these results indicate that CteG is important for centrosome amplification which in turn is essential for normal *C.t.* replication in primary cells.

### CETN2 is essential for centrosome amplification during *C.t.* infection

Our data indicate that CteG is required for centrosome amplification during chlamydial infection, which we hypothesize is due to its interaction with CETN2. To determine if CETN2 is integral for centrosome amplification during chlamydial infection, we used siRNA to knockdown CETN2 expression. Due to low abundance of the CETN2 protein, we were unable to detect it by Western blotting even in standard HeLa cell lysates without immunoprecipitation. Thus, we used Quantigene to determine knockdown efficiency, achieving an average of a 12-fold decrease in CETN2 mRNA transcript. We found that knockdown of CETN2 resulted in a significant decrease in the percentage of cells with supernumerary centrosomes, a trend that was noted in both WT L2 or *cteG::aadA* infected cells. In alignment with our data showing a significant decrease in the percentage of cells with supernumerary centrosomes between WT L2 and *cteG::aadA* infected cells (Fig. 6), we saw a significant difference between WT L2 and *cteG::aadA* infected cells in the control KD. But, there was not a significant difference in the CETN2 KD condition between WT L2 and *cteG::aadA* (Fig. 6). However, the exacerbated decrease in cells with supernumerary centrosomes in the CETN2 KD cells infected with WT L2 or *cteG::aadA* further supports a role for the CteG-CETN2 interaction in centrosome amplification, implicating the necessity of both CteG and CETN2 for this phenotype.

**Fig. 6.**
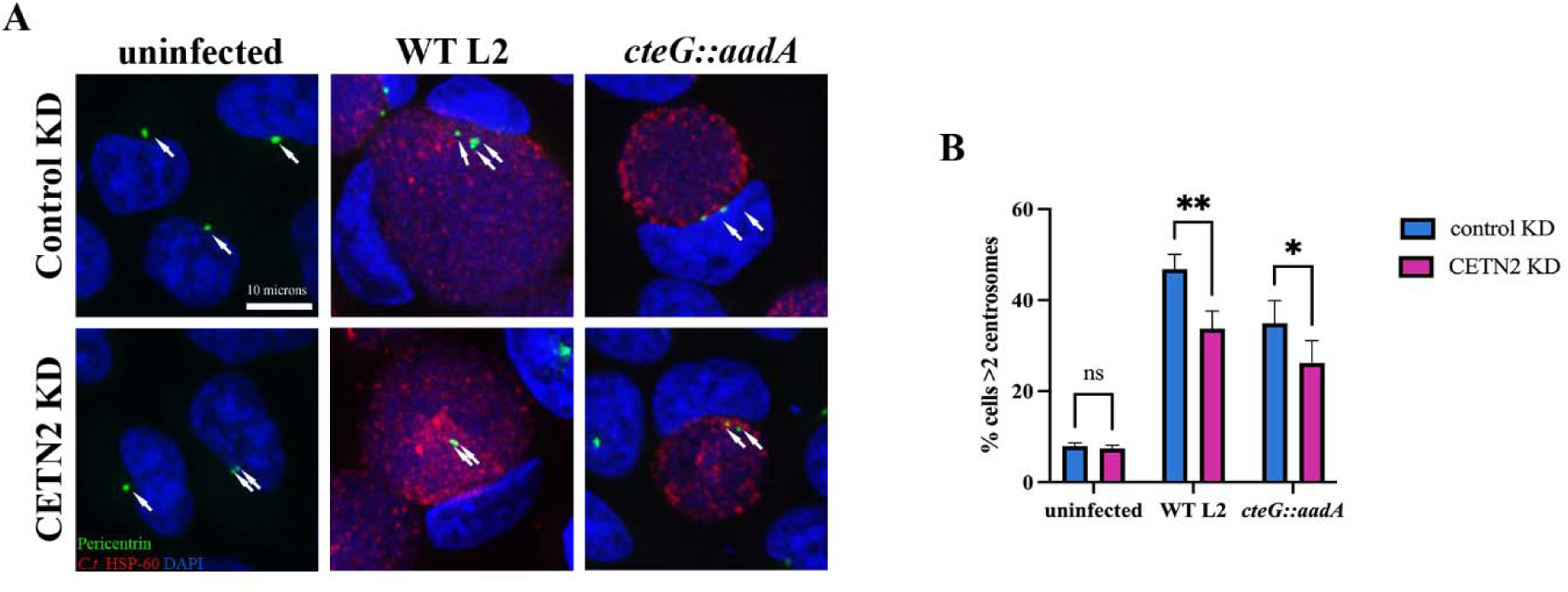
CETN2 is necessary for CteG-mediated centrosome amplification. (*A*) Representative images of HeLa cells depleted (CETN2 KD, *right*) or not (Control KD, *left*) of CETN2 and infected with WT L2 or *cteG::aadA* for 36 hours. Cells were stained with *C.t.* HSP-60 (red), pericentrin (green), and DAPI (blue). White arrows indicate centrosomes (Scale bar, 10 microns). (*B*) Quantification of cells with supernumerary centrosomes (>2) at 36 hours post infection in CETN2 KD and control KD cells. Error bars are SD, * P< 0.05, ** P<0.01. Significance was determined using two-way ANOVA followed Tukey’s multiple comparisons test. Data are representative of 2 replicates.

## Discussion

As an obligate intracellular pathogen, *C.t.,* from the confines of its inclusion, must engage several host organelles and signaling pathways to carve out its unique replicative niche. To achieve these feats, *C.t.* releases an arsenal of cT3SS effector proteins into the host cell, the function of most remains largely unknown. Our data indicates that CteG, through interactions with centrin-2, induces centrosome amplification during chlamydial infection (Fig. 7). CteG represents the first bacterial factor to target centrin proteins and notably, our findings begin to dissect how a bacterial pathogen induces such cellular abnormalities as centrosome amplification, that have canonically been associated with viral infections.

**Fig. 7.**
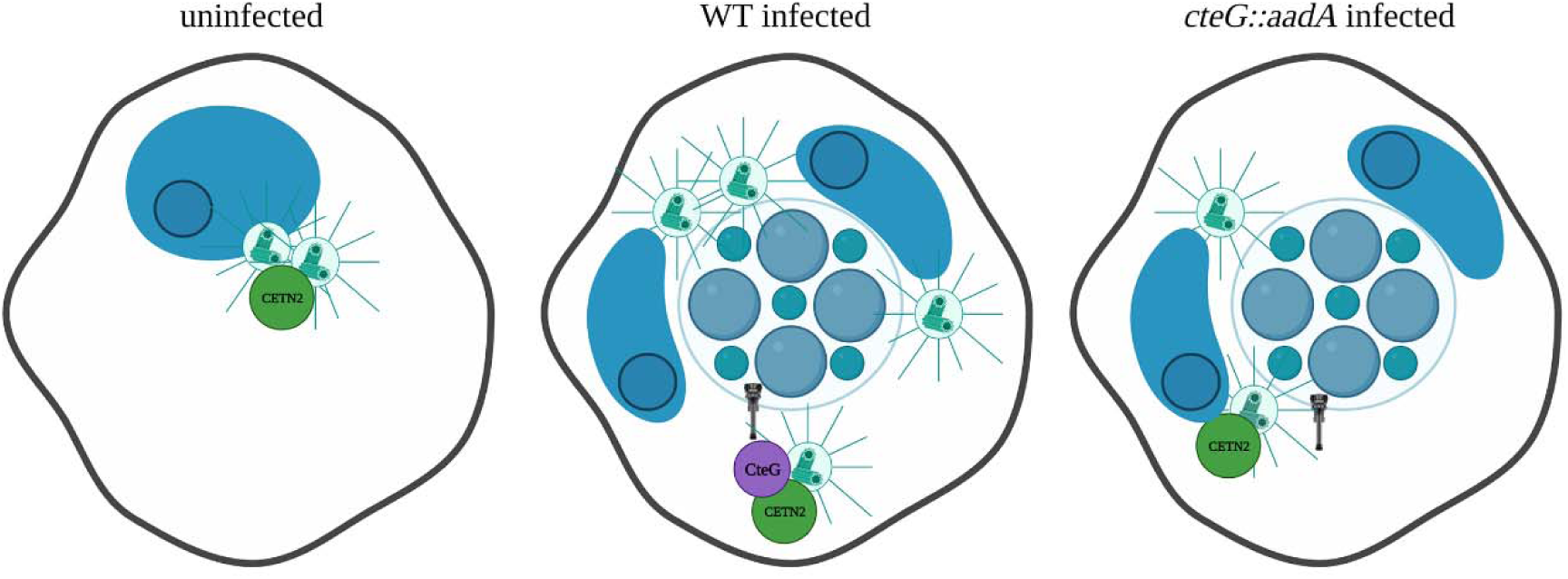
Working model. Uninfected cells maintain a normal number and localization of centrosomes. In chlamydia infected cells, supernumerary centrosomes, induced through CteG-CETN2 interactions, help promote inclusion anchoring at the microtubule-organizing center. In the absence of CteG or CETN2, centrosome amplification is decreased.

Our work highlights the importance of CETN2 in the regulation of centrosome amplification and further provides useful insight into how centrosome amplification may be regulated. CETN2 is an important structural component of centrosomes and is a key regulator of centriole duplication (32). As a member of the EF-hand superfamily, it harbors distinct helix-loop helix domains that coordinate calcium binding (33, 34, 36). Binding of calcium is presumed to be important for target recognition with low-affinity sites becoming higher-affinity sites in the presence of calcium (37). While CETN2 possesses four EF-hand domains, the important calcium-regulatory sites for human centrin proteins appears to be the pair of EF hands at the C-terminus (38). Our data indicate that an intact calcium binding domain of EF hand 4 is important for CteG binding (Fig. 3E). Given the importance of calcium binding for target recognition, we predict that calcium binding to EF hand 4 induces a conformational change that enables CteG binding. As centrosome assembly in mammalian cells requires CETN2 association with other proteins or protein complexes including CaM (calmodulin) and CP110 (39), hSfi1 (40, 41), and hPOC5 (42) for appropriate centrosome duplication and mitotic spindle assembly, how CteG binding impacts these associations warrants further study. As many of these interactions occur at the C-terminus of CETN2, binding of CteG to this region may obscure CETN2’s interaction with other host proteins impairing regulation of the centrosome duplication process, suggesting CteG is acting as an agonist to promote centrosomes amplification.

Our findings add to the growing body of literature that link *C.t.* infection to induction of gross cellular abnormalities, such as supernumerary centrosomes, mitotic spindle defects, multinucleation, aneuploidy, and blocked cytokinesis that were initially described over 15 years ago, but are still mechanistically undefined (17–21). To date, most studies have been performed in HeLa cells or E6/E7 transformed cell lines, clouding whether observed phenotypes are due to *C.t.* infection or are artifacts of HPV infection in these cell lines. Recent work by Wang et al. showed that centrosome amplification is an additive effect between HPV and *C.t*. (18), but this occurs through different mechanisms. Using HPV-negative cell lines, they show that centrosome amplification requires progression through the cell cycle and may result from a cytokinesis defect. Building on these findings, our new data indicate that centrosome amplification can also be induced through CteG-CETN2 interactions. Intriguingly, HeLa or A2EN cells infected with the *cteG::aadA* have significantly reduced centrosomes relative to cells infected with WT L2, yet the number of centrosomes present in the *cteG::aadA* infected cells are still elevated relative to uninfected cells (Fig. 5B). Strikingly though, in primary cells, the percent of *cteG::aadA* infected cells with supernumerary centrosomes mirrored that of uninfected cells (Fig. 5C) suggesting that transformed cell lines inherently synergize with chlamydial infection to promote supernumerary centrosomes. This could be due to the transforming factors themselves (E6/E7 or other oncogenes) or something inherent to immortalized cells that promotes centrosome amplification in conjunction with chlamydial infection. Thus, our new data, in conjunction with previous studies indicate that *C.t.* may employ multiple methods to drive centrosome amplification during infection. In addition to failed cytokinesis, which could lead to supernumerary centrosomes, previous studies also suggest that the secreted factor CPAF may also be important for centrosome amplification (21). Regardless of the method, our data show for the first time that centrosome amplification is important for chlamydial replication and inclusion development. In primary cells, we observe a growth defect for the *cteG::aadA* strain, which is notably absent in the immortalized HeLa and A2EN cells. We postulate that centrosome amplification, driven by HPV E6/E7 proteins in the immortalized line, partially compensates for the inability of *C.t.* to cause supernumerary centrosome formation in the absence of CteG. Alternatively, other changes caused by HPV E6/E7 may be involved. While it is clear that *C.t.* needs elevated centrosomes for normal replication and inclusion development, why they are needed remains unknown.

While our data clearly indicate that CteG-CETN2 interactions are necessary for centrosome amplification, no difference in centrosome clustering or positioning was noted. A recent study by Sherry et al. revealed that the inclusion membrane protein, Dre1, interacts with dynactin to reposition host organelles, namely centrosomes, to help with the positioning of the *C.t.* inclusion at the MTOC (43). Dre1 is responsible for overriding normal host centrosome clustering mechanisms to allow *C.t.* to position centrosomes in close proximity to the inclusion. Other Incs, including CT223/IPAM and CT288 bind to centrosome components (22, 44, 45). IPAM has been associated with centrosome amplification and failed cytokinesis in IPAM-transfected cells (22). IPAM also recruits CEP170, a centrosomal protein, to control microtubule organization and assembly from the inclusion (45). CT288 was shown to interact with human centrosomal protein CCDC146 and is partially responsible for recruiting it to the inclusion membrane during infection, potentially playing a role in inclusion anchoring at the MTOC (44). Collectively, these studies support a role for Inc proteins in the positioning of the inclusion at the MTOC. Thus, we hypothesize that CteG is responsible for the initial amplification of centrosomes, and then Incs become involved for repositioning centrosomes and microtubules within the host to aid in the positioning of the inclusion at the MTOC.

Previous work on CteG showed localization to the Golgi or plasma membrane depending on the stage of the infection (25). More recent work implicated CteG in *C.t.* lytic exit from the host (46). Mota et al. found decreased host cell cytotoxicity in *cteG*::*bla* infected cells, indicating a role for this effector in host cell lysis at the end of the *C.t.* lifecycle to facilitate release of infectious chlamydia. We hypothesize that centrosome amplification is necessary for helping localize the inclusion to the MTOC, which would be necessary in early and mid-stages of the infection cycle. Centrosomes are less clustered in *C.t.* infected cells (43), so CteG may be localizing with these centrosomes around the mature inclusion along the plasma membrane, where it could then help facilitate lytic exit later in the infection cycle. As centrosomes serve as important microbial tracks, it is possible that less microtubules encompass the inclusion in a CteG mutant strain, leading to changes in lytic exit.

Taken together, we propose a model where upon infection, CteG is secreted and interacts with CETN2 to induce centrosome amplification to aid in the positioning of the inclusion at the MTOC with the help of inclusion membrane proteins (Fig. 7). We speculate that changes in cell cycle progression (as measured by EdU staining) are a downstream effect of CteG’s primary effect on centrosome amplification, as this amplification process likely slows down the host cell cycle, and centrosome duplication is heavily linked to cell cycle progression. Further characterization of the CteG-CETN2 interaction is necessary to understand the mechanistic underpinnings of this interaction and how it leads to centrosome amplification. This would contribute to our understanding of how *C.t.* induces gross host cell abnormalities that are also hallmarks of cancer, potentially providing a link between *C.t.* infection and increased cancer risk.

## Materials and Methods

### Bacterial and Cell Culture

*C.t.* serovar L2 (LGV 434/Bu) was propagated in HeLa 229 cells (American Type Tissue Culture), and EBs were purified using a gastrograffin density gradient as previously described (47). HeLa cells were grown in RPMI 1640 with L-Glutamine (Thermo Fisher Scientific) supplemented with 10% Fetal Bovine Serum (Gibco), sodium bicarbonate, sodium pyruvate, and gentamicin at 37°C with 5% CO_2_. A2EN cells (Kerafast) were propagated in keratinocyte-serum free media (K-SFM) (Thermo Fisher Scientific) supplemented with 0.16 ng/ml epidermal growth factor (EGF), 25 μg/ml bovine pituitary extract (BPE), 0.4 mM CaCl_2_, and gentamicin (48, 49). Primary cervical cells were derived from normal HPV-negative cervical tissue obtained through the University of Iowa Tissue Procurement Core from a consenting donor who underwent a hysterectomy for endometriosis (IRB#201103721 and IRB#199910006). Normal cervical epithelial cells were isolated as previously described (50) and were maintained in K-SFM (Thermo Fisher Scientific) without CaCl_2_.

### Cloning

TargeTronics was used to predict TargeTron insertion sites for CT144. gBlocks were obtained from Integrated DNA Technologies (Table S1) and were cloned into the HindIII/BsrGI site of pACT (51).

### Chlamydia Transformation

*C.t.* EBs were transformed as previously described (52) with minor modifications. Briefly, fresh *C.t.* lysates were mixed with 5 µg plasmid DNA and 10 µl of 5X transformation mix (50 mM Tris pH 7.4 and 250 mM CaCl_2_) in a total volume of 50 µl. Mixtures were incubated at room temperature for 30 min, resuspended in RPMI, and applied to 2-wells of a 6-well plate of confluent HeLa cells. Plates were centrifuged at 900 x g for 30 min.

At 18 hours post-infection, 0.3 µg/ml penicillin G was added. Infectious progenies were harvested every 48 h and used to infect a new HeLa cell monolayer until viable inclusions were present (∼2-3 passages). Expression of FLAG-tagged proteins was confirmed by western blotting. For TargeTron mutants, successful insertion into the target gene was confirmed by PCR.

### Yeast Suppressor Screen

To identify putative suppressors of CteG toxicity, a yeast suppressor screen was carried out as previously described (28, 53). Briefly, CteG was cloned into the KpnI/XbaI site of pYesNTA and the resulting plasmid (pYesNTA-CteG) was transformed into *S. cerevisiae* W303. To assess toxicity, transformants were serially diluted and spotted onto uracil dropout medium containing glucose or galactose as the sole carbon source. To identify yeast ORFs that suppress CteG toxicity, the pYEp13 genomic library (ATCC no. 37323) was transformed into the W303-CteG strain. Transformants were plated on uracil leucine dropout medium containing galactose. From a total transformation of ∼1.0X10^5^, we obtained 69 colonies. Plasmids were isolated from clones that consistently suppressed the toxicity of CteG and isolated plasmids were retransformed into W303-CteG to confirm suppression. To identify yeast ORFs present, suppressor plasmids were sequenced using pYEp13 seq F and pYEp13 seq R (Table S1). Sequences were analyzed using the yeast genome database (https://www. yeastgenome.org/). To validate suppression, putative suppressors were then individually cloned into p415-ADH (33).

### Affinity Purification

HeLa cells were infected at an MOI of 2 with *C.t.* strains expressing a FLAG-tagged effector protein, under tetracycline inducing conditions (10 ng/ml) for 24 hours. Cells were lysed in eukaryotic lysis solution (ELS) (50 mM Tris HCl, pH 7.4, 150 mM NaCl, 1 mM EDTA, and 1% Triton-X 100) and spun at 12,000 x g for 20 min. Supernatants were incubated with 60µl preclearing beads (mouse IgG agarose, Millipore Sigma) for 2 hours. The precleared lysate was incubated with 30µl FLAG beads (anti-FLAG M2 Affinity Gel, Millipore Sigma) overnight. The beads were washed 6 times with ELS without detergent. For mass spectrometry, samples were stored in 50 mM ammonium bicarbonate prior to digestion and analysis. For western blotting, proteins were eluted from the beads in NuPAGE LDS Sample Buffer (Thermo Fisher Scientific) and boiled for 5 minutes.

### Mass Spectrometry

Beads containing samples were washed with 25mM ammonium bicarbonate and digested with 0.5 µg trypsin (Pierce, Thermo Fisher Scientific, MS Grade) using a CEM microwave reactor for 30 minutes at 55°C. Digested peptides were extracted twice using 50% acetonitrile plus 5% formic acid, lyophilized to dry, and resuspended in 5% acetonitrile plus 0.1% formic acid. For LC/MS, samples were injected into an UltiMate 3000 UHPLC system coupled online to a high resolution Thermo Orbitrap Fusion Tribrid mass spectrometer. Peptides were separated by reversed-phase chromatography using a 25 cm Acclaim PepMap 100 C18 column with mobile phases of 0.1% formic acid and 0.1% formic acid in acetonitrile; a linear gradient from 4% formic acid in acetonitrile to 35% formic acid in acetonitrile over the course of 45 minutes was employed for peptide separation. The mass spectrometer was operated in a data dependent manner, in which precursor scans from 300 to 1500 m/z (120,000 resolution) were followed by collision induced dissociation of the most abundant precursors over a maximum cycle time of 3 seconds (35% NCE, 1.6 m/z isolation window, 60 s dynamic exclusion window). Raw LC-MS/MS data was searched against a database containing UniProt_Human and Chlamydia_trachomatis_L2434Bu using Mascot 2.8. Tryptic digestion was specified with a maximum of two missed cleavages, while peptide and fragment mass tolerances were set to 10 ppm and 0.6, respectively. Quantitation was done using Mascot Average method using Mascot Distiller 2.8.2.

### Co-immunoprecipitations

Co-immunoprecipitations were performed on either co-transfected HeLa cells or cells that were transfected using Lipofectamine LTX (Thermo Fisher Scientific) and subsequently infected at an MOI of 2.5 for 24 hours. Cells were lysed with ELS and spun at 12,000 x g for 20 min. Supernatants were incubated with 50 µl FLAG magnetic beads (Pierce^™^ Anti-DYKDDDDK, Thermo Fisher Scientific) for 2 hours. The beads were washed 6 times with ELS without detergent. Proteins were eluted from the beads in NuPAGE LDS Sample Buffer (Thermo Fisher Scientific) and boiled for 5 minutes prior to analysis by Western blotting.

### Western blotting

Samples were separated by SDS-PAGE and transferred to PVDF membranes. Blots were blocked in 5% milk in Tris-buffered saline with Tween 20 (TBST). Membranes were probed with an anti-GFP (Novus), anti-FLAG (Thermo Fisher Scientific), or anti-HA (Millipore Sigma) primary antibody and goat anti-rabbit HRP conjugate (BioRad) secondary antibody. Results were collected from at least 3 independent experiments.

### Immunofluorescence

HeLa cells were co-transfected with CETN2-dsRed and GFP-tagged *C.t.* effectors CteG, TmeA, or empty vector. Cells were fixed in 4% formaldehyde, permeabilized with 0.1% Triton-X and stained with DAPI. Images were taken on a Nikon Eclipse Ti2 microscope. Images were analyzed for colocalization using Fiji Coloc2 function to calculate a Pearson’s R value. Values greater than 0.7 are considered significant.

### Centrosome Staining

Immunofluorescence centrosome staining was done as previously established with modification (18, 43). HeLa, A2EN, or primary cervical cells were infected with the appropriate strains of *C.t.* at an MOI of 1 by centrifugation at 700 x g for 30 min. At 36 hours post-infection, cells were fixed on ice with cold methanol for six minutes and blocked for two hours at room temperature in 0.1% Triton-X in PBS with 2% FBS. Cells were stained with anti-pericentrin (abcam) and anti-Chlamydia HSP60 (Millipore Sigma). Dylight-488 and Dylight-594 (Thermo Fisher Scientific) secondaries were used along with DAPI (Thermo Fisher Scientific) to stain the nuclei. Images were captured using a Leica DFC7000T confocal microscope equipped with Leica software. At least 10 images were collected per coverslip, with three technical replicates per biological replicate, with at least 2 biological replicates.

### Centrosome measurements

For centrosome number measurements, maximal projection images obtained from confocal imagery were used for counting the number of centrosomes per cell. Cells with >2 centrosomes were considered to have “supernumerary centrosomes.” All centrosomes of infected cells from at least 10 images per technical replicates were counted, with at least 2 biological replicates per cell type. To measure centrosome clustering, Fiji was used to create a polygon encompassing all centrosomes in a cell and the area of this shape was measured. To measure centrosome spread, the distance from each centrosome to the nearest edge of the nucleus was determined. Centrosomes on the nucleus were given a value of zero. A total of 100 measurements were taken for each condition. For infected conditions, only *C.t.* infected cells were analyzed.

### Edu Labeling

Confluent HeLa cell monolayers were infected with the appropriate strains of *C.t.* at an MOI of 1 by centrifugation at 700 x g for 30 min. At 36 hours post-infection, cells were incubated with 10 µM EdU for 30 minutes at 37°C using the Click-iT EdU Cell Proliferation kit (Thermo Fisher Scientific, C10337). Samples were fixed with 4% formaldehyde and permeabilized with 0.5% Triton-X. At least 10 images were collected (by Nikon Eclipse Ti2 microscope) per coverslip, with three technical replicates per biological replicate, with at least 2 biological replicates.

### Growth Curve

HeLa cells were infected at an MOI of 2.5 on ice. After 30 mins, media was changed, and plates were moved to 37°C with 5% CO_2_ to stimulate bacterial uptake. At 0 or 48h, cells were lysed in water and lysates were used to infect fresh monolayers of HeLa cells. Titer plates were fixed with methanol 24 h post-infection and stained with anti-chlamydial LPS (Novus). IFUs at 48 hours were normalized to the WT L2 IFUs at 0 hours.

### siRNA Knockdown

HeLa cells were transfected using Dharmafect with SmartPool siRNA for CETN2 or ON-TARGET*plus* Cyclophilin B control according to manufacturer’s protocol (Dharmacon). At 36 hours post-transfection, cells were infected with the appropriate strains of *C.t.* at an MOI of 1 by centrifugation at 700 x g for 30 minutes and incubated for 36 hours. Cells were fixed and stained for centrosomes as described above. Knockdown efficiency was determined using QuantiGene^™^ (Thermo Fisher Scientific) according to the manufacturer’s protocol.

### Statistics

When necessary, statistical analysis was performed using GraphPad Prism 9.3.0 software. One-way and two-way ANOVA’s were used followed by Tukey’s multiple comparisons with p<0.05 (*), p<0.01 (**), and p<0.001 (***).

## Acknowledgements

We thank Drs. Derek Fisher and Luís Jaime Mota for sharing the *cteG::aadA* strain. We also thank Drs. Joanne Engel and Jessica Sherry for assistance with troubleshooting centrosome staining and measurements. Additionally, we would like to thank Paige McCaslin for technical support, Isaias Herring for assistance with bacterial titering, and Jabeena CA for critical review of this manuscript. We acknowledge grant support from the NIH (B.S. T32 AI007511, M.M.W. R01 AI150812, R01 AI155434).

