## Supplementary material for "The *Chlamydia trachomatis* type III secreted effector protein CteG induces centrosome amplification through interactions with centrin-2": Sup Fig. 1-3 and Table

**Supporting Information**

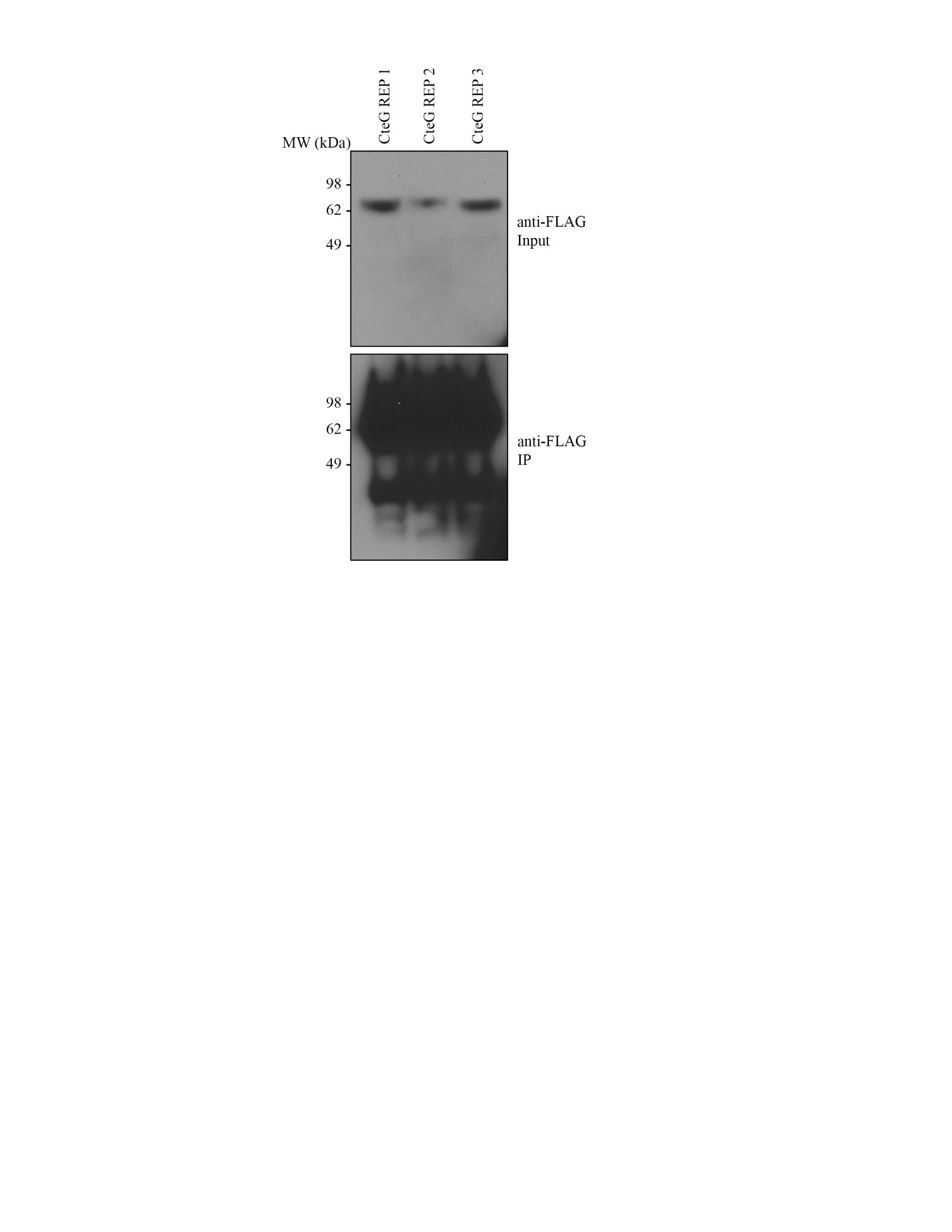

**Fig. S1.** Check of CteG-FLAG expression for AP-MS samples. IP of FLAG-tagged CteG from HeLa cells infected with *C.t.* expressing FLAG-tagged CteG. A portion of the three replicates sent for mass spectrometry.

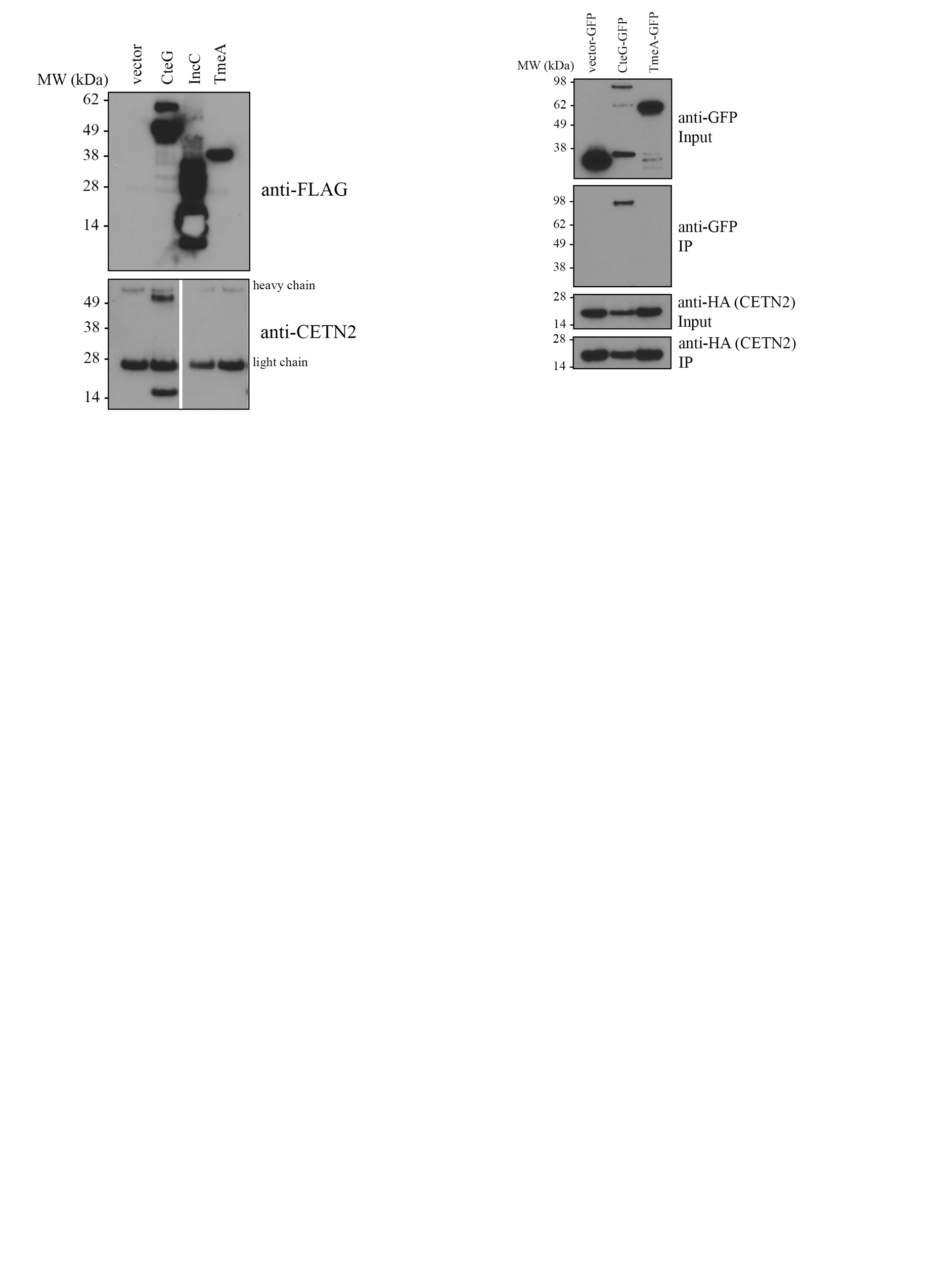

**Fig. S2.** CteG co-IPs with endogenous CETN2. Co-IP of FLAG-tagged effectors from HeLa cells infected with *C.t.* expressing the FLAG-tagged vector, CteG, IncC, or TmeA. Blot probed with antibody against CETN2 to look for endogenous interaction with *C.t.* effectors.

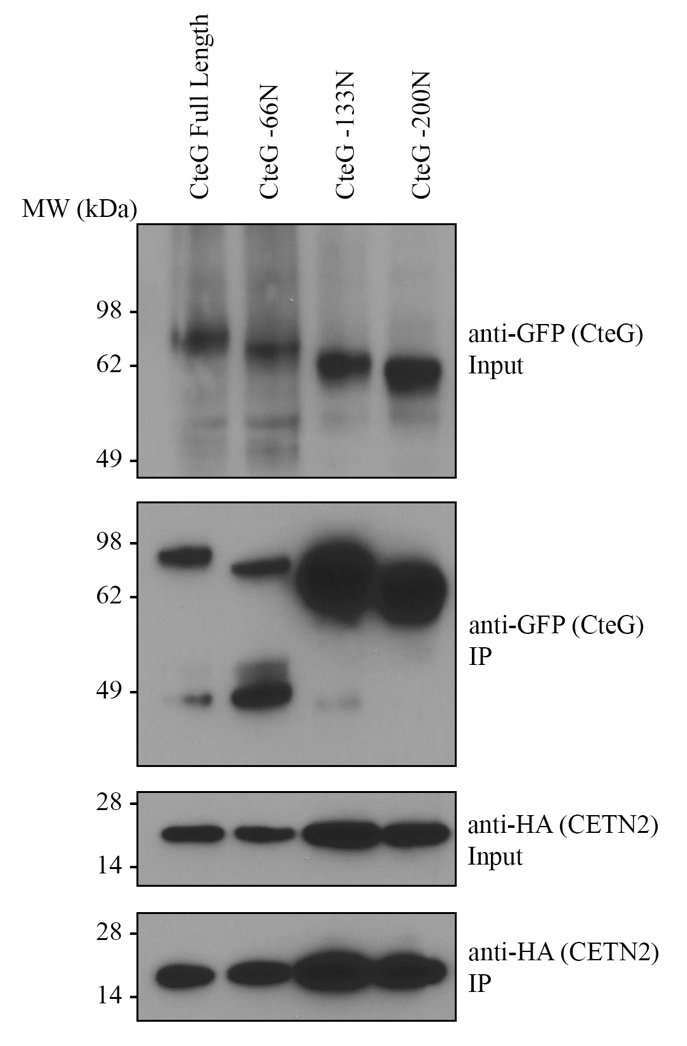

**Fig. S3.** N-terminus is dispensable for CteG interaction with CETN2. Co-IP of HA-tagged CETN2 from HeLa cells co-transfected with GFP-tagged CteG truncations and CETN2-HA.

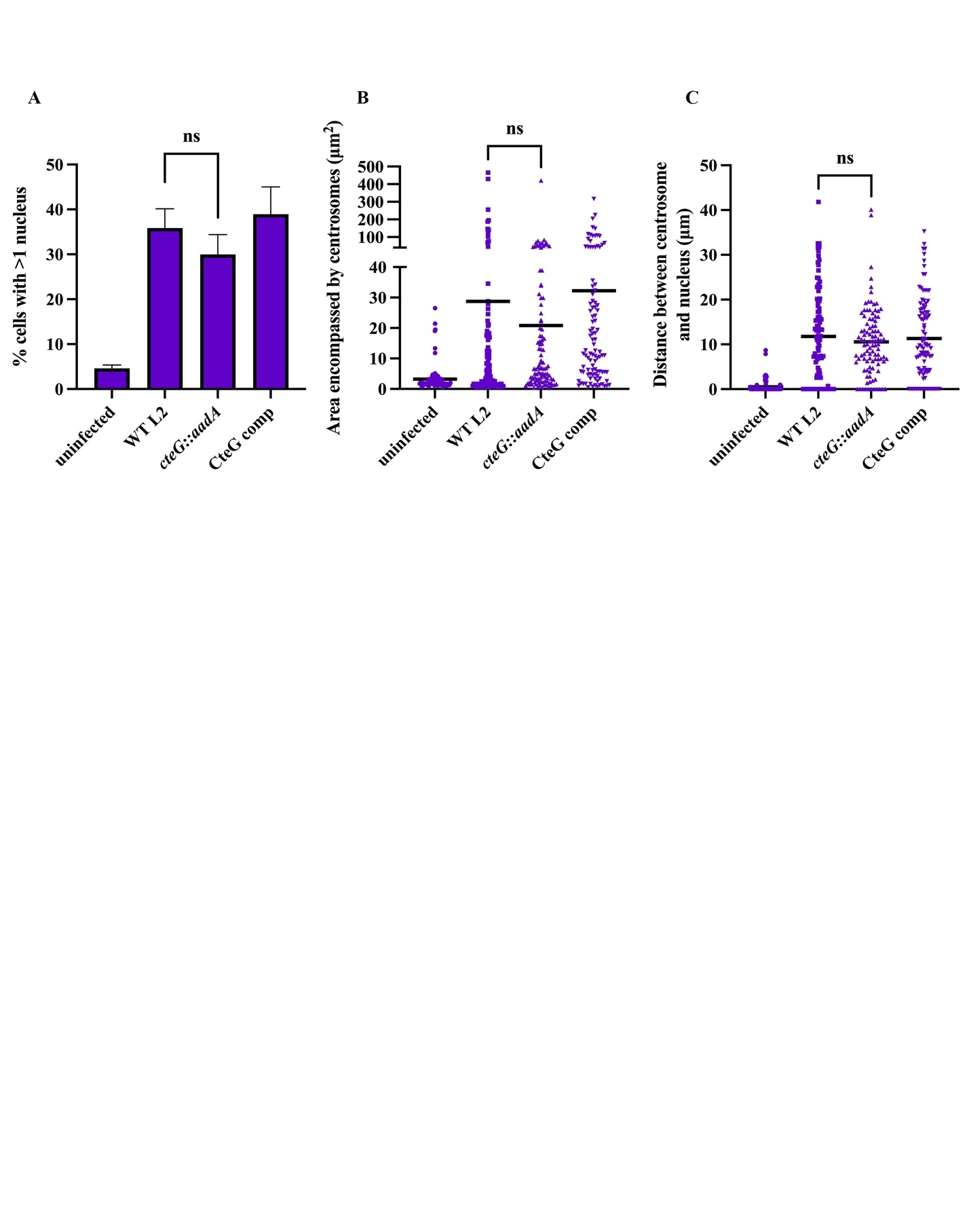

**Fig. S4.** CteG does not affect multinucleation, centrosome clustering, or centrosome distribution. (*A*) Quantification of cells with multinucleation (>1) at 36 hours post infection in A2EN cells represented as percent of total cells. Error bars are SD. Data are representative of 2 replicates. (*B*) Measurement of the area encompasses by centrosomes in A2EN cells. Black bars represent the mean. Data are representative of 100 counted cells. (*C*) Measurement of the distance between centrosomes to the nearest edge of the nucleus in A2EN cells. Black bars represent the mean. Data are representative of 100 counted cells. (*A-C*) Significance was determined using one-way ANOVA followed by Tukey’s multiple comparisons test.

**Table S1.** Primers used in this study.

| **Name** | **Use** | **Sequence** |
| --- | --- | --- |
| CT144 33/34s | TargeTron mutagenesis | TTCCCCTCTAGAAAAAAGCTTATAATTATCCTTAACTATCGATGTTGTGCGCCCAGATAGGGTGTTAAGTCAAGTAGTTTAAGGTACTACTCTGTAAGATAACACAGAAAACAGCCAACCTAACCGAAAAGCGAAAGCTGATACGGGAACAGAGCACGGTTGGAAAGCGATGAGTTACCTAAAGACAATCGGGTACGACTGAGTCGCAATGTTAATCAGATATAAGGTATAAGTTGTGTTTACTGAACGCAAGTTTCTAATTTCGATTATAGTTCGATAGAGGAAAGTGTCTGAAACCTCTAGTACAAAGAAAGGTAAGTTAGAAACATCGACTTATCTGTTATCACCACATTTGTACAATCTG |
| LtrB F | TargeTron mutagenesis | TTCCCCTCTAGAAAAAAGCTTATAATTATCCTTA |
| LtrB R | TargeTron mutagenesis | CAGATTGTACAAATGTGGTGATAACAGATAAGTC |
| pYEp13 sequencing F | Yeast suppressor screen sequencing | ACTACGCGATCATGGCGA |
| pYEp13 sequencing R | Yeast suppressor screen sequencing | TGATGCCGGCCACGATGC |
| APC2 HindIII F | Yeast suppressor screen | CCAAGCTTATGTCATTTCAGATTACCCCAAC |
| APC2 SalI R | Yeast suppressor screen | CCGTCGACTCATGAGTTTTTATGCCCATTTT |
| CETN2 KpnI F | CETN2 truncations | CCGGTACCATGGCCTCCAACTTTAAGAAGGC |
| CETN2 XhoI R | CETN2 truncations | CCCTCGAGTTAATAGAGGCTGGTCTTTTTCAT |
| CETN2 1-400 XhoI R | CETN2 truncations | CCCTCGAGTTACCAACTCCTTGGCCACGC |
| CETN2 1-300 XhoI R | CETN2 truncations | CCCTCGAGTTATTTCTCAGACATTTTCTGGGTCA |
| CETN2 1-447 XhoI R | CETN2 truncations | CCCTCGAGTTAAGCTTCATCAATCATTTCCTGC |
| CETN2 1-486 XhoI R | CETN2 truncations | CCCTCGAGTTAGAACTCTTGCTCACTGACCTCTC |
| CETN2 99-516 KpnI F | CETN2 truncations | CCGGTACCCGGGAAGCTTTTGATCTTTTCG |
| CETN2 198-516 KpnI F | CETN2 truncations | CCGGTACCGAAGAAATTAAGAAAATGATAAG |
| CteG +1 KpnI F | Yeast suppressor screen | CCGGTACCAATGTCATTTGGTATTGGTAGTGCTT |
| CteG Xba1 R | Yeast suppressor screen/Ectopic expression | CCTCTAGACTAGATAGAGGAGCTTTGCACAC |
| CteG KpnI F | Ectopic Expression | CCGGTACCATGTCATTTGGTATTGGTAGTGC |
| CteG NotI F | CteG truncations/ pBomb4 plasmid cloning | CCGCGGCCGCATGTCATTTGGTATTGGTAGTGCTT |
| CteG FLAG KpnI R | CteG truncations | CCGGTACCTTACTTATCGTCGTCATCCTTGTAATCGATAGAGGAGCTTTGCACACCT |
| CteG FLAG SalI R | CteG truncations/ pBomb4 plasmid cloning | CCGTCGACTTACTTATCGTCGTCATCCTTGTAATCGATAGAGGAGCTTTGCACACCT |
| CteG -198N KpnI F | CteG truncations | CCGGTACCCCAATGGTAGGGACGTACTCAG |
| CteG -399N KpnI F | CteG truncations | CCGGTACCTCTGCAAGAGGTGCTGGTTCC |
| CteG -600N KpnI F | CteG truncations | CCGGTACCTTTCTTGCTTTAGGAGGAT |
| CteG -198C Xba1 R | CteG truncations | CCTCTAGAGGATGATTGCAAATGCGTTAAAC |
| CteG -399C Xba1 R | CteG truncations | CCTCTAGACTTACAAAGAAGCATGGTCAACT |
| CteG -600C Xba1 R | CteG truncations | CCTCTAGAACCGCCTGCTAATCCGCTG |
| CteG -51C FLAG SalI R | pBomb4 plasmid cloning | CCGTCGACTTACTTATCGTCGTCATCCTTGTAATCTTCTCTCATAAGATCTTTACTTT |
| CteG -99C FLAG SalI R | pBomb4 plasmid cloning | CCGTCGACTTACTTATCGTCGTCATCCTTGTAATCCTCTAATAGGGATAGGAAAGAAC |
| CteG -150C FLAG SalI R | pBomb4 plasmid cloning | CCGTCGACTTACTTATCGTCGTCATCCTTGTAATCACGCACTTCTGCTCGAGTTCTGT |
| TmeA KpnI F | Ectopic expression | CCGGTACCATGAGTATTCGACCTACTAATGGGAG |
| TmeA NotI R | Ectopic expression | CCGCGGCCGCGTCTAAGAAAACAGAAGAAGTT |
| TmeA KpnI +1 F | Yeast suppressor screen | CCGGTACCAATGAGTATTCGACCTACTAATGGGAG |
| TmeA Xbal R | Yeast suppressor screen | CCTCTAGATTAGTCTAAGAAAACAGAAGAAGTTATGAC |
| TmeA Not F | pBomb4 plasmid cloning | CCGCGGCCGCATGAGTATTCGACCTACTAATGGGAG |
| TmeA Flag SalI R | pBomb4 plasmid cloning | CCGTCGACTTACTTATCGTCGTCATCCTTGTAATCGTCTAAGAAAACAGAAGAAGTTATGAC |
| IncC NotI F | pBomb4 plasmid cloning | CCGCGGCCGCATGACGTACTCTATGTCCGATA |
| IncC FLAG SalI R | pBomb4 plasmid cloning | CCGTCGACTTACTTATCGTCGTCATCCTTGTAATCGCTTACATATAAAGTTTGAGGAT |

**Table S2.** Complete list of filtered AP-MS peptides for CteG.

| **Protein Description** | **Accession** | **Database** | **Avg. Score** | **Mass** | **Avg. Num. of Sig. Matches** | **Avg. Num. of Sig. Sequences** |
| --- | --- | --- | --- | --- | --- | --- |
| Actin, cytoplasmic 2 | P63261 | UniProt_  Human | 2445 | 41766 | 77 | 20 |
| CdsQ, CT672/CTL0041 | A0A654L4P8 | Chlamydia_  trachomatis_L2434Bu | 1049 | 41172 | 28 | 15 |
| Putative RNA-binding protein Luc7-like 2 | Q9Y383 | UniProt_  Human | 681 | 46486 | 20 | 10 |
| Putative RNA-binding protein Luc7-like 1 | Q9NQ29 | UniProt_  Human | 122 | 43701 | 3 | 2 |
| Centrin-2 | P41208 | UniProt_  Human | 844 | 19726 | 21 | 9 |
| CteG, CT105/CTL0360 | A0A654L6L6 | Chlamydia_  trachomatis_L2434Bu | 425 | 68204 | 11 | 7 |
| Filamin-A | Q60FE5 | UniProt_  Human | 493 | 278053 | 20 | 16 |
| Cleavage and polyadenylation specificity factor subunit 6 | F8WJN3 | UniProt_  Human | 456 | 52238 | 7 | 4 |
| Tropomodulin-3 | Q9NYL9 | UniProt_  Human | 428 | 39570 | 13 | 10 |
| Keratin, type I cytoskeletal 9 | P35527 | UniProt_  Human | 252 | 62027 | 4 | 4 |
| Girdin | A0A2R8Y7B1 | UniProt_  Human | 243 | 185366 | 9 | 7 |
| 4F2 cell-surface antigen heavy chain | J3KPF3 | UniProt_  Human | 186 | 68059 | 5 | 4 |
| Signal recognition particle receptor subunit beta (Fragment) | H7C4H2 | UniProt_  Human | 157 | 17602 | 4 | 4 |
| 50S ribosomal protein L16, rplP, CT521/CTL00783 | A0A654L7G0 | Chlamydia_  trachomatis_L2434Bu | 134 | 15765 | 3 | 3 |
| Transaldolase, CT313/CTL0565 | A0A654L6E7 | Chlamydia_  trachomatis_L2434Bu | 252 | 36138 | 5 | 4 |
| Peptidyl-prolyl cis-trans isomerase A | P62937 | UniProt_  Human | 110 | 18001 | 4 | 3 |
| 60S acidic ribosomal protein P2 | P05387 | UniProt_  Human | 141 | 11658 | 4 | 4 |
| DnaK, CT396/CTL0652 | A0A654L6Q8 | Chlamydia_  trachomatis_L2434Bu | 147 | 70800 | 4 | 4 |
| ATP synthase subunit alpha, mitochondrial (Fragment) | K7EK77 | UniProt_  Human | 131 | 22186 | 6 | 6 |
| Adenosylmethionine decarboxylase | A0A5F9ZHD5 | UniProt_  Human | 111 | 53138 | 3 | 3 |
| Calmodulin-1 | P0DP23 | UniProt_  Human | 143 | 16827 | 3 | 3 |
| Pre-mRNA 3'-end-processing factor FIP1 (Fragment) | H0Y8P7 | UniProt_  Human | 116 | 29476 | 2 | 2 |
| Activated RNA polymerase II transcriptional coactivator p15 | P53999 | UniProt_  Human | 173 | 14386 | 7 | 6 |
| IncG, CT118/CTL0373 | A0A654LGL6 | Chlamydia_  trachomatis_L2434Bu | 98 | 17529 | 2 | 1 |
| Catenin (Cadherin-associated protein), alpha 1, isoform CRA_a | G3XAM7 | UniProt_  Human | 125 | 92663 | 2 | 2 |
| Importin-9 | Q96P70 | UniProt_  Human | 91 | 115889 | 4 | 2 |
| Peptidyl-prolyl cis-trans isomerase B | P23284 | UniProt_  Human | 312 | 23728 | 11 | 8 |
| Ribosomal protein L19 | J3KTE4 | UniProt_  Human | 154 | 23233 | 4 | 3 |
| GrpE, CT395/CTL0651 | A0A654L6N2 | Chlamydia_  trachomatis_L2434Bu | 87 | 21655 | 3 | 2 |
| Mediator of RNA polymerase II transcription subunit 1 | Q15648 | UniProt_  Human | 181 | 168373 | 3 | 2 |
| Brain acid soluble protein 1 | P80723 | UniProt_  Human | 99 | 22680 | 2 | 2 |
| Serine/threonine-protein phosphatase 6 regulatory subunit 1 | Q9UPN7 | UniProt_  Human | 107 | 96664 | 2 | 2 |
| Spectrin beta chain | A0A087WUZ3 | UniProt_  Human | 75 | 274659 | 3 | 2 |
| Albumin | A0A087WWT3 | UniProt_  Human | 94 | 45118 | 2 | 1 |
| Ras-related protein Rab-5C | P51148 | UniProt_  Human | 89 | 23468 | 2 | 1 |
| Protein disulfide-isomerase A3 | P30101 | UniProt_  Human | 102 | 56747 | 4 | 4 |
| DnaJ, CT341/CTL0595 | A0A654L6M8 | Chlamydia_  trachomatis_L2434Bu | 93 | 41890 | 3 | 3 |
| 60S ribosomal protein L7a (Fragment) | Q5T8U3 | UniProt_  Human | 92 | 21531 | 3 | 3 |
| Importin subunit beta-1 (Fragment) | J3QR48 | UniProt_  Human | 78 | 16388 | 2 | 2 |
| S-adenosylmethionine synthase isoform type-2 | P31153 | UniProt_  Human | 129 | 43633 | 2 | 2 |
| 40S ribosomal protein S6 | A2A3R5 | UniProt_  Human | 85 | 24953 | 3 | 3 |
| Cofilin, non-muscle isoform (Fragment) | E9PP50 | UniProt_  Human | 102 | 17766 | 3 | 3 |
| HLA class I histocompatibility antigen, A alpha chain | A0A0G2JPD3 | UniProt_  Human | 97 | 44397 | 2 | 2 |
| CADD, CT610/CTL0874 | A0A654L7A9 | Chlamydia_  trachomatis_L2434Bu | 73 | 26816 | 2 | 2 |
| CdsJ, CT559/CTL0822 | A0A654L770 | Chlamydia_  trachomatis_L2434Bu | 73 | 35489 | 2 | 1 |
| Peptidyl-prolyl cis-trans isomerase-like 4 | Q8WUA2 | UniProt_  Human | 59 | 57189 | 3 | 2 |
| FACT complex subunit SPT16 | Q9Y5B9 | UniProt_  Human | 88 | 119838 | 4 | 4 |
| Histone H1.2 | P16403 | UniProt_  Human | 75 | 21352 | 2 | 2 |
| Heat shock 70 kDa protein 1B | A0A0G2JIW1 | UniProt_  Human | 89 | 70066 | 3 | 3 |
| FACT complex subunit SSRP1 | Q08945 | UniProt_  Human | 69 | 81024 | 2 | 2 |
| Chaperonin GroEL, CT110/CTL0365 | A0A654L5L1 | Chlamydia_  trachomatis_L2434Bu | 62 | 58054 | 3 | 3 |
| 50S ribosomal protein L17, RplQ, CT506/CTL0768 | A0A654L6Q2 | Chlamydia_  trachomatis_L2434Bu | 59 | 16142 | 2 | 2 |
| FolP, CT613/CTL0877 | A0A654L9I1 | Chlamydia_  trachomatis_L2434Bu | 80 | 50230 | 2 | 2 |
| 60S ribosomal protein L28 | H0YKD8 | UniProt_  Human | 57 | 19060 | 3 | 2 |
| Spindlin interactor and repressor of chromatin-binding protein | Q9BUA3 | UniProt_  Human | 73 | 41011 | 2 | 2 |
| 60S ribosomal protein L34 | P49207 | UniProt_  Human | 44 | 13284 | 2 | 1 |
| 60 kDa heat shock protein, mitochondrial | P10809 | UniProt_  Human | 39 | 61016 | 2 | 2 |
| 50S ribosomal protein L13, RplM, CT125/CTL0380 | A0A654L5X5 | Chlamydia_  trachomatis_L2434Bu | 54 | 16839 | 2 | 2 |
| 60S ribosomal protein L23a (Fragment) | H7BY10 | UniProt_  Human | 86 | 17792 | 3 | 2 |
| Membrane-associated progesterone receptor component 1 | O00264 | UniProt_  Human | 68 | 21658 | 2 | 2 |
| 40S ribosomal protein S12 | P25398 | UniProt_  Human | 105 | 14505 | 2 | 1 |
| Centrin-3 | E5RJF8 | UniProt_  Human | 59 | 22409 | 2 | 2 |
| 30S ribosomal protein S9, RpsI, CT126/CTL0381 | A0A654L6N5 | Chlamydia_  trachomatis_L2434Bu | 56 | 14533 | 2 | 2 |
| Vimentin | P08670 | UniProt_  Human | 58 | 53619 | 2 | 2 |
| 60S ribosomal protein L13 | P26373 | UniProt_  Human | 81 | 24247 | 3 | 3 |
| 60S ribosomal protein L5 (Fragment) | A0A2R8Y6J3 | UniProt_  Human | 42 | 27028 | 2 | 2 |
| Calcium homeostasis endoplasmic reticulum protein | J3QK89 | UniProt_  Human | 61 | 104868 | 2 | 1 |
| ATP synthase subunit d, mitochondrial | F5H608 | UniProt_  Human | 55 | 8910 | 2 | 2 |
| eIF-2-alpha kinase activator GCN1 | Q92616 | UniProt_  Human | 41 | 292572 | 2 | 2 |
| 30S ribosomal protein S10, RpsJ, CT436, CTL0695 | A0A654L6R0 | Chlamydia_  trachomatis_L2434Bu | 61 | 11862 | 2 | 2 |
| Pre-mRNA-splicing factor 38A | Q8NAV1 | UniProt_  Human | 44 | 37453 | 2 | 2 |
| Uncharacterized *C.t.* protein CT504/CTL0766 | A0A654LIJ4 | Chlamydia_  trachomatis_L2434Bu | 52 | 32014 | 2 | 2 |
| Importin-8 | F5H815 | UniProt_  Human | 26 | 3990 | 2 | 2 |
| 60S ribosomal protein L21 | P46778 | UniProt_  Human | 34 | 18553 | 2 | 2 |
| 60S ribosomal protein L26 | J3KSS0 | UniProt_  Human | 31 | 7570 | 2 | 1 |
| 50S ribosomal protein L21, RpIU, CT420/CTL0677 | A0A654L7I1 | Chlamydia_  trachomatis_L2434Bu | 62 | 12155 | 2 | 2 |
